# Comparative Analysis of Lipid Nanoparticles in Pfizer-BioNTech and Moderna COVID-19 Vaccines: Insights from Molecular Dynamics Simulations

**DOI:** 10.1101/2024.10.04.616619

**Authors:** Malay Ranjan Biswal, Sudip Roy, Jayant K Singh

## Abstract

COVID-19 vaccines, such as Pfizer-BioNTech’s BNT162b and Moderna’s mRNA-1273, have demonstrated robust efficacy. However, direct comparisons of their delivery vehicles remain limited. Notably, BNT162b requires storage at -80°C, while mRNA-1273 is stored at -20°C. This discrepancy in storage temperatures may be influenced by differences in the structure and stability of the lipid nanoparticles (LNPs) used in these vaccines. Ionizable lipids, such as SM-102 in Moderna’s vaccine and ALC-0315 in Pfizer’s vaccine, play a crucial role in LNP stability and function, affecting endosomal escape, cellular uptake, and drug release. Understanding these variations is essential for optimizing vaccine delivery systems. In our study, we use molecular dynamics simulations with the coarse-grained Martini forcefield to compare the LNPs in Moderna and Pfizer’s COVID-19 vaccines, providing insights at an experimental scale. Our findings indicate that the ionizable lipid tail of BNT162b (ALC-0315) exhibits a higher degree of branching, resulting in a more bifurcated appearance compared to the structure of the ionizable lipids in mRNA-1273 (SM-102).

## Introduction

Vaccines, from Edward Jenner’s pioneering smallpox vaccine to cutting-edge innovations, revolutionized public health by preventing diseases and saving lives. Traditional vaccines, which contain antigens from pathogens or artificially created mimics of pathogen components, stimulate immune responses, protecting specific pathogens.^1,2^ However, these conventional vaccines present many challenges, including safety concerns, storage requirements, and the need for multiple administrations. ^3,4^

mRNA vaccines, developed by scientist Katalin Kariko, represent a significant advancement in vaccine technology. These vaccines offer benefits such as rapid development,^5,6^ precise antigen expression,^7,8^ and reduced toxicity.^9^ Besides their application as COVID-19 vaccines, mRNAs against multiple diseases like cancer, ^10–12^ mRNA encoding immunomodulatory factors, ^13^ tumour suppressor genes, ^14,15^ and antibodies^16^ are either in preclinical or clinical phases. Despite many appealing features of mRNA as vaccine or drugs, there are multiple challenges in mRNA delivery, primary because of the instability of mRNA due to enzymatic degradation by ribonucleases present in the blood.^17^ Also, the highly negative charge and large size create insufficient intracellular delivery.

There are many methods developed to overcome these challenges. One of them is the delivery of mRNA via vesicles called liposomes or lipid nanoparticles (LNPs).^18–20^ These vesicles carry not only RNAs but several other drugs and cellular materials like proteins, lipids and DNA, from source to distant sites via biofluids. The vesicles protect nucleic acids and other biological molecules from degradation *in vivo* and deliver them to the targeting sites without interfering with immune activation. These liposomes and nanoparticles earlier have been used in the treatment of cancer^21^ and vaccines.^22,23^ And many of these liposomes have been used as a carrier in clinical trials to deliver anticancer, anti-inflammatory, antibiotic, antifungal, anesthetic, and other drugs and gene therapies, as they help the biomolecules to cross the negative potential cell membrane barrier. These LNPs effectively deliver nucleic acids by forming micelles or vesicles to encapsulate drugs. Recently, two authorized COVID-19 vaccines, mRNA-1273^24,25^ and BNT162b^26^ use LNP to deliver antigen mRNA.

The efficacy of COVID-19 vaccines, such as Pfizer-BioNTech’s BNT162b and Moderna’s mRNA-1273, has been well-established, though limited direct comparisons between their delivery vehicle exist. Existing comparisons predominantly focus on mRNA performance or in vivo studies,^27,28^ overlooking a detailed examination of the LNP structure and stability, a critical aspect of these vaccines. Notably, the presence of ionizable lipids, such as SM-102 in Moderna’s vaccine and ALC-0315 in Pfizer’s vaccine, along with other compositional differences, can significantly influence the overall structure and stability of LNPs. ^29^ Moreover, they play important role in endosomal escape, cellular uptake and drug release. Understanding these variations is crucial for optimizing vaccine delivery systems and enhancing our comprehension of their performance and stability. This gap in research emphasizes the need for a comprehensive investigation into the structural implications of specific lipid components on LNPs, shedding light on potential avenues for further vaccine refinement. In this work we will be comparing the structure and stability of LNPs of Moderna and Pfizer’s COVID-19 vaccine using coarse-grained (CG) molecular dynamics study with CG Martini forcefield. This will provide insights at a larger scale comparable to experimental size.

## Modelling and computational details

In this study, we investigated the formation and stability of vesicles derived from LNPs used in the Moderna and Pfizer COVID-19 vaccines. Coarse-grained molecular dynamics simulations were employed for this purpose. To perform these simulations, we required the coarse-grained structures of RNA and individual lipid molecules. These structures were then utilized to create vesicles, conduct simulations, and analyze the results.

### Initial configuration

#### mRNA secondary structure

Obtaining accurate secondary structures for RNA molecules for MD simulations presents several challenges due to the inherent complex characteristics of RNA. Addressing these challenges requires specialized tools, with ContextFold playing a crucial role. ContextFold^30^ employs a comparative modeling approach that integrates experimental data, sequence information, and structural constraints to predict 3D structures for RNA.

In our research, we employed ContextFold to predict the secondary structure of a specific mRNA sequence. This sequence encodes the spike protein found in the SARS-CoV-2 virus (the virus responsible for COVID-19). The entire mRNA sequence is quite long—about 3,822 nucleotides. However, simulating such a large sequence would require a very large system size, which could be challenging. To address this, we focused on a smaller part of the mRNA—the one that codes for the spike domain of the virus. This domain is crucial because it interacts with the ACE2 receptor in human cells. Due to limitations in the prediction tool, we analyzed 500 nucleotides at a time. The predicted structure from ContextFold served as the starting point for more detailed molecular dynamics (MD) simulations and further analysis.

#### Creating vescicle

In the development of lipid nanoparticles (LNPs) for the Moderna and Pfizer COVID-19 vaccines, specific lipid compositions were employed. Moderna LNPs utilized the ionizable lipid SM-102 (heptadecan-9-yl 8-((2-hydroxyethyl)(6-oxo-6-(undecyloxy)hexyl)amino) octanoate), phospholipid DPPC (1,2-Dipalmitoylphosphatidylcholine), PEGylated lipid PEG2000-DMG, and cholesterol in a ratio of 50:10:38.5:1.5. Similarly, for Pfizer LNPs, the ionizable lipid ALC-0315 ((4-hydroxybutyl)azanediyl)bis(hexane-6,1-diyl)bis(2-hexyldecanoate), DPPC, PEGylated lipid ALC-0159, and cholesterol were combined in a ratio of 46.3:9.4:42.7:1.6. The lipid compositions were sourced from Verbeke et al.’s study.^31^ The composition of all the lipids and mRNAs are given in the table 1.

**Table 1:** Lipid composition in the vesicles of Moderna and Pfizer.

|  | mRNA-1273 |  | BNT162b |  |
| --- | --- | --- | --- | --- |
|  | #Molecules | %Weight | #Molecules | %Weight |
| ILIP | 4450 | 6.664 | 3806 | 6.874 |
| DPPC | 3444 | 5.157 | 3878 | 5.253 |
| CHOL | 142 | 0.095 | 120 | 0.073 |
| PEG | 1016 | 2.282 | 1134 | 2.176 |
| mRNA | 2 | 0.553 | 2 | 0.500 |
| Water | 682144 | 85.124 | 753077 | 85.010 |
| Na | 998 | 0.124 | 998 | 0.113 |

In our study, we employed the MARTINI coarse-grained (CG) version 2 for molecular dynamics simulations. Following the Martini philosophy, we represented four heavy atoms as a single CG bead (referred to as “four-to-one”). Additionally, we implemented three-to-one and two-to-one mappings for smaller beads when necessary. The initial structures of ionizable lipids and PEG molecules were created and optimized using Avogadro. Subsequently, we coarse-grained these molecules according to the CG-Martini guidelines. The bead type for these lipids were given in supplementary figures, S1-S4. For other lipids, such as DPPC and cholesterol, we obtained their coarse-grained structures from the Martini 2 database. The all atom structure of the mRNA obtained from ContextFold was coarse grained using the “martinize-nucleotide python script”. ^32^

To construct vesicles mimicking these LNPs, bilayers were first generated using PACKMOL.^33^ PACKMOL helps create initial configuration for MD simulation by packing molecule in defined region of space. Coarse-grained structure of lipids and water were placed within a simulation box dimensions of 10 nm *×* 10 nm *×* 10 nm based on the corresponding lipid ratios mentioned above. Subsequently, 100 ns of MD simulations were conducted to equilibrate the system. Following equilibration, the simulation box was replicated four times in both the X and Y directions, resulting in the creation of a large bilayer, which is 16 times larger than the starting bilayer configuration.

The BumPy tool^34^ was then employed to transform the large bilayer into a vesicle with an approximate diameter of 40 nanometers. The BumPy tool helps in the creation of lipid bilayers of varying curvature and composition which can be used for the starting configuration of MD simultion. Before employing BumPy tool, the PEG molecules present in the lower leaflet of the bilayer were removed. Approximately 750000 water beads including 10% water beads and required amount of sodium ions were added. This newly formed vesicle underwent an additional 100 ns equilibration through MD simulations. Subsequently, two mRNAs generated using the ContextFold tool were introduced into the vesicle. Finally, a comprehensive NPT MD simulation, lasting 1500 ns, was conducted to explore the dynamic behavior and stability of the vesicle structure containing the mRNAs. The methods was illustrated in the figure 1.

**Figure 1:**
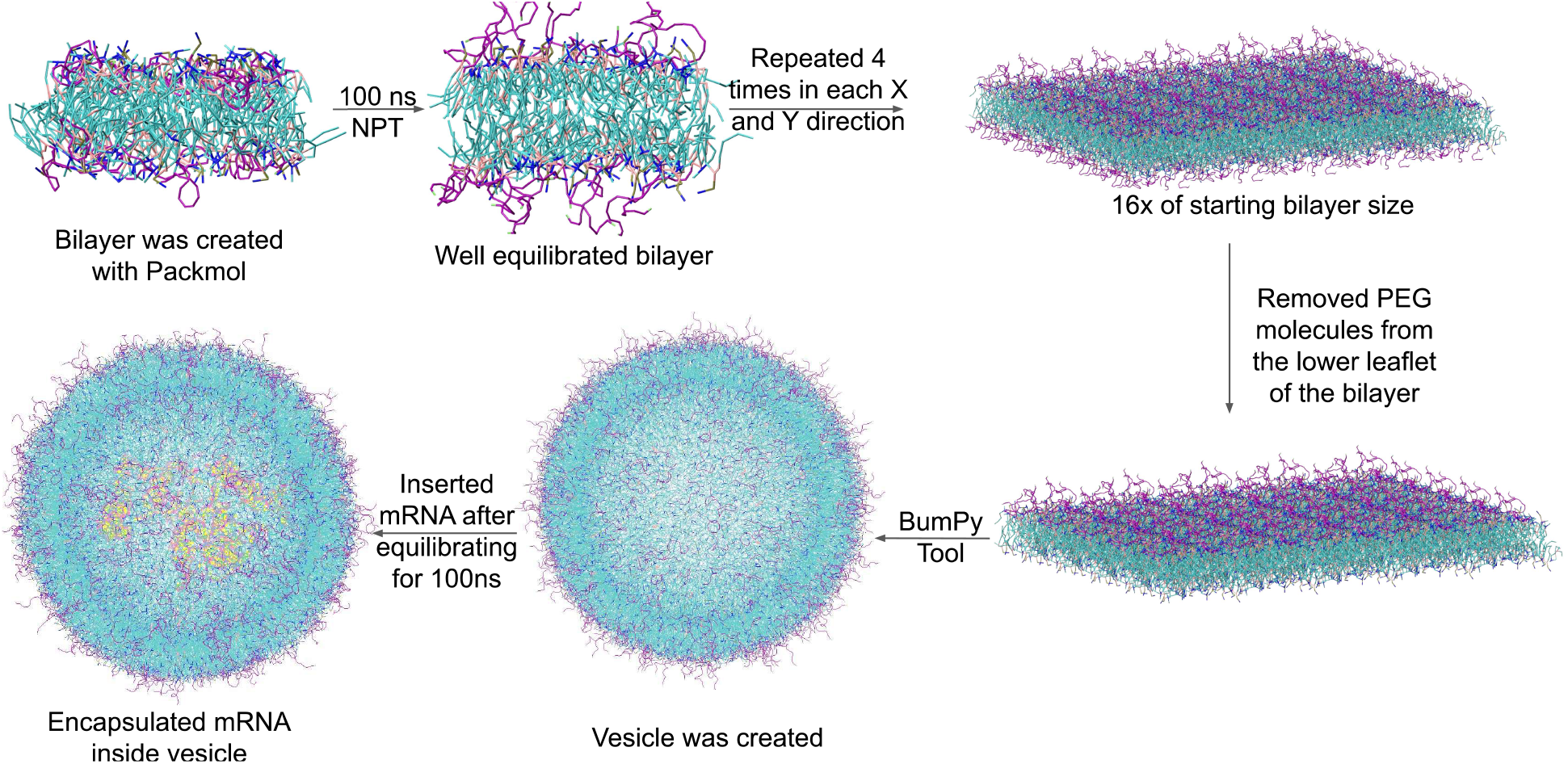
Methods followed in creating vesicles.

### Computational details

All molecular dynamics simulations were conducted utilizing GROMACS version 2020.^35^ The system was minimised with steepest descent minimization. A Verlet cut-off scheme was employed, incorporating a short-range electrostatic cut-off of 1.2 nm and a short-range van der Waals cut-off of 1.2 nm. Neighbor searching utilized a Verlet cut-off scheme with a frequency set to update the neighbor list every 10 steps. The relative dielectric constant was set at 15. Temperature coupling was achieved through velocity rescaling, with a reference temperature of 310 K for physiological temperature and a coupling time constant of 0.1. For NPT simulation, an isotropic Parrinello-Rehman pressure coupling of 1 bar was applied, with a time constant of 4 ps. Periodic boundary conditions were maintained in all directions.

## Results and discussion

In our study, we created two lipid vesicles using a composition similar to the LNPs found in the Moderna and Pfizer COVID-19 vaccines, as mentioned in the methods section. Our goal was to explore the structure and stability of these two vesicles. To do this, we closely examined the structure and arrangement of lipids within the vesicle bilayers.

### Formation of stable lipid nanoparticles

To explore the assembly of the liposomal model and assess the formation of LNP from Moderna and Pfizer, the Radial Distribution Function (RDF) parameter was employed. RDFs were calculated by selecting specific atoms, such as the head group of ionizable lipids or phospholipids, as reference points. Subsequently, the distance and distribution of neighboring atoms in relation to the reference atoms were measured.

In Figure 2, the RDF index is plotted along the vertical axis, while the horizontal axis depicts the distance of ionizable and phospholipid atoms from the reference atom. In both the lipids, the RDF shows the DPPC lipids are closer to each other and aggregated. This behavior is seen as prominent in BNT162b compared to mRNA-1273. The RDF graph reveals a well-ordered ionizable and phospholipid atoms arrangement, indicating favorable van der Waals interactions. The RDF parameter shows that both ionizable and phospholipids are organized and stable, indicating that the 1500 ns time scale is suitable for forming and stabilizing the liposomal model structure.

**Figure 2:**
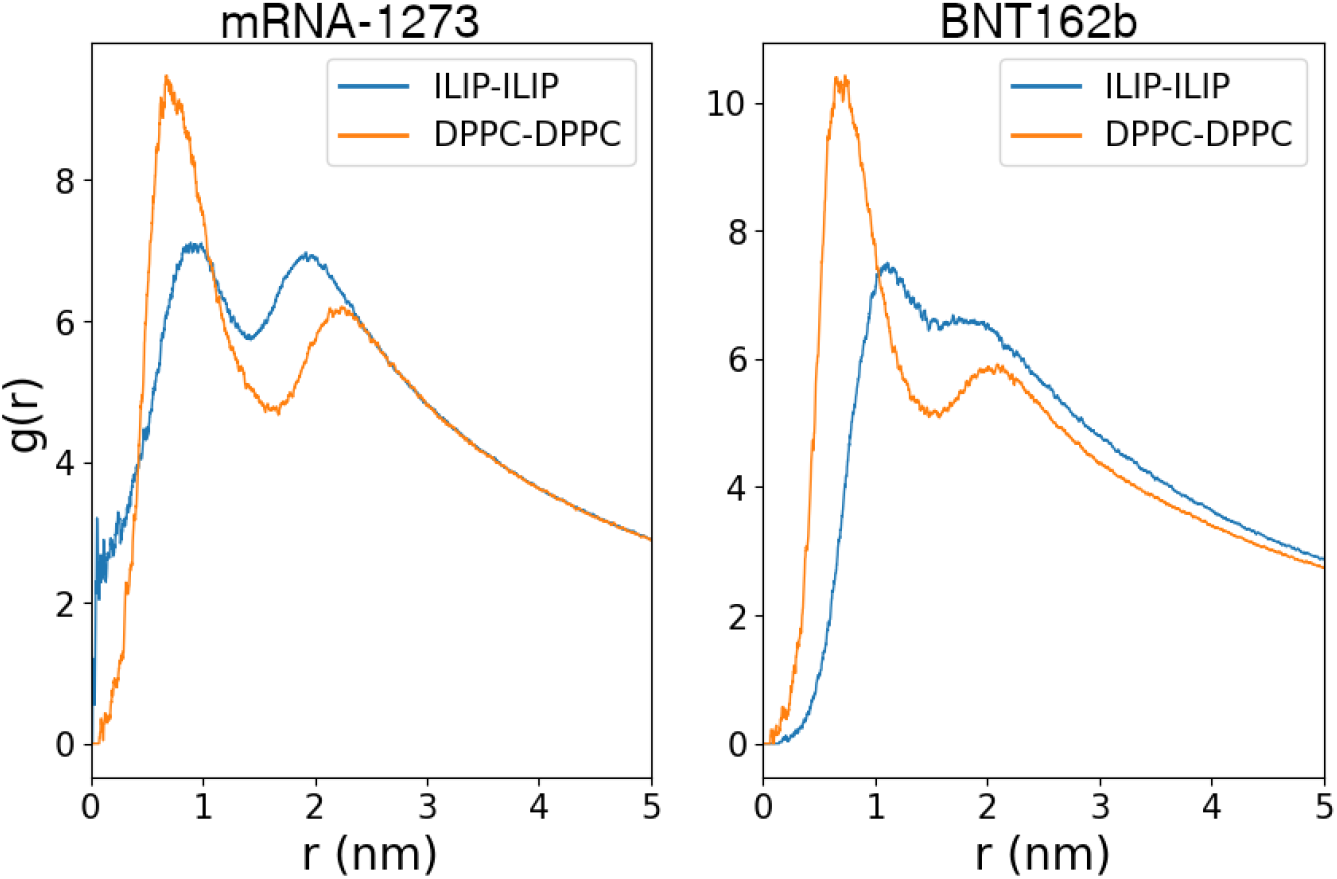
The Radial distribution function between ionizable lipids (ILIP-ILIP) and phospholipids (DPPC-DPPC). Lipids in mRNA-1273

To understand the arrangement of phospholipids and ionizable lipids we looked at the different section of the vesicles. The cross section of vesicles were taken at difference regions from bottom to the middle as shown in the Figure S5. Corresponding section of mRNA-1273 and BNT162b is shown in Figure 3, to understand the lipid arrangement. We observed distinct lipid arrangements within the vesicles containing mRNA-1273 lipids compared to those with BNT162b lipids. The key distinction lies in the ionizable lipid composition. Specifically, in the case of mRNA-1273, the ionizable lipid SM-102 possesses a single branched tail out of its two tails. Conversely, the ionizable lipids of BNT162b, represented by ALC-0315, feature branched tails on both sides. This structural variation significantly impacts the lipid packing within the vesicles. Lipids in mRNA-1273 exhibit a more densely packed and organized lipid arrangement. The presence of a single branched tail in SM-102 allows for tighter packing, whereas the dual branched tails in ALC-0315 lead to a more dispersed distribution.

**Figure 3:**
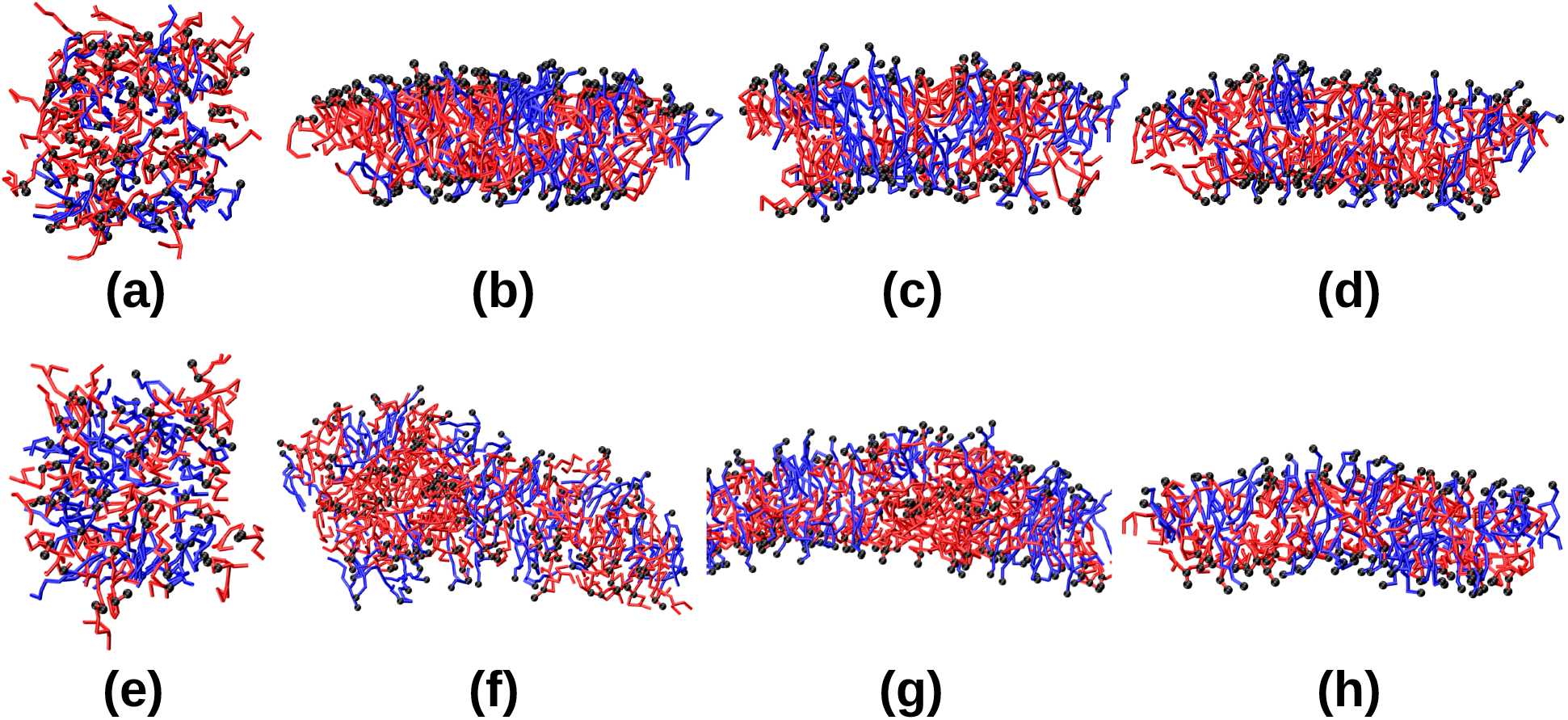
The distribution of DPPC and ionizable lipids at different cross section of the vesicle. It is seen that the arrange of lipids in the (a)-(d) mRNA-1273 is more ordered compared to BNT162b (e)-(h). DPPC is shown in blue and ionizable lipids are shown in red. Cholesterol, PEG molecules, mRNA and Water molecules are not shown for clarity. The region of vesicle which has been shown here is shown in supplementary figures S7 and S8.

The radial density of head group of the phospholipids and ionizable lipids of both mRNA-1273 and BNT162b were calculated and plotted in the figure 4. The densities were calculated with respect to the center of vesicle by considering the center as center of geometry of all the lipid molecules. It is observed that the densities of the head group have very sharp peaks, almost 40% higher than BNT162b, indicating that the lipids are well arranged in the spherical vesicle and formed a very clear distinct bilayer. In comparison to mRNA-1273, the lipids in the BNT162b exhibit lower densities and broader profiles.

**Figure 4:**
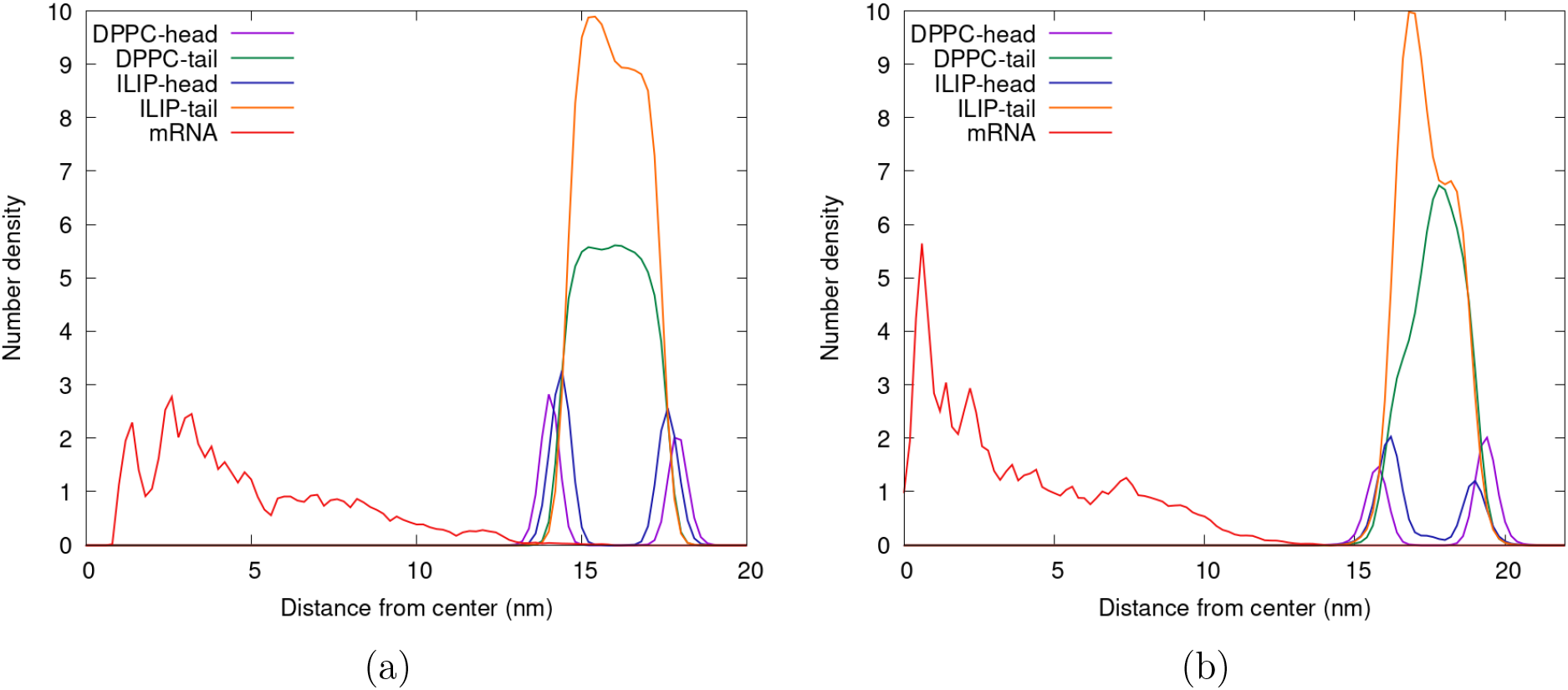
Radial density of the head group and tail group of DPPC and ionizable lipid (ILIP) in (a) mRNA-1273 and (b) BNT162b. The lipids in the mRNA-1273 are more dense and better arranged compared to the lipids in BNT162b.

### Lipid stability

The stability and formation of colloidal particles, specifically LNPs, are governed by a combination of attractive van der Waals and repulsive electrostatic interactions, as outlined by the Derjaguin– Landau– Verwey– Overbeek (DLVO) theory.^36^ Phospholipids initially bond via hydrogen, later stabilized by van der Waals interactions. Despite being individually weak, these interactions play a vital role in shaping and stabilizing lipid-based structures like liposomes. They promote lipid clustering and organization within the membrane, facilitated by both electrostatic and van der Waals forces. The figure 5 illustrates two important interactions, the coulombic and van der Waals interactions, between lipids in LNPs from mRNA-1273 and BNT162b, contribute to vesicle formation and stability. Although both contribute to overall stability, van der Waals interactions play a significant role than electrostatic interactions. While the difference in the van der Waals interaction is negligible, the electrostatic interaction is almost 20% higher in BNT162b lipids.

**Figure 5:**
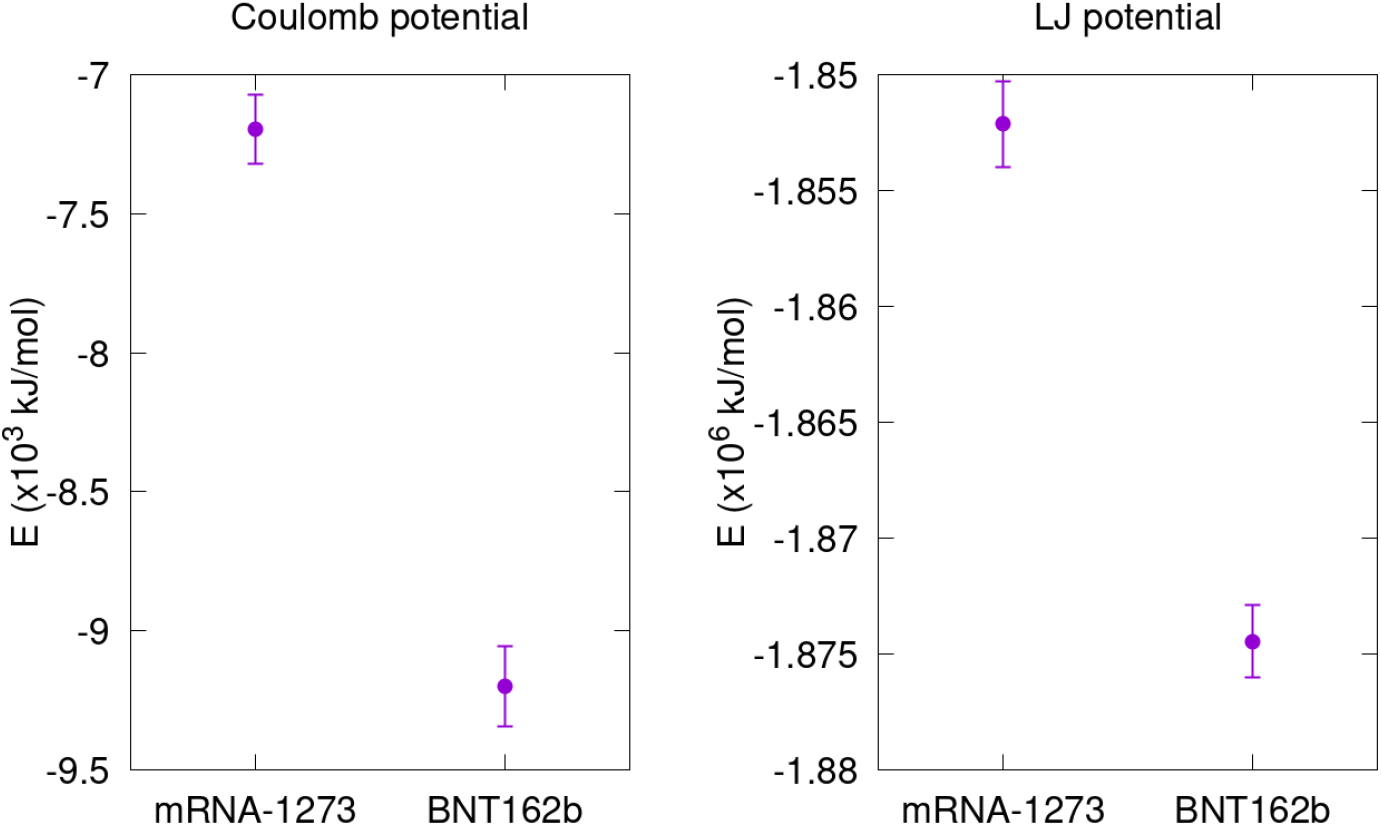
Interaction energy between all the lipid molecules of mRNA-1273 and BNT162b, shows that both the lipids are very stable with minimal difference in the Coulomb and LJ interaction.

We also determined the bilayer thickness of the vesicles by measuring the difference between the outer and inner diameters of the LNPs multiple times and averaging the values. The thickness of the mRNA-1273 LNPs was approximately 3.8 nm, while it was approximately 3.1 nm in BNT162b. It’s important to note that this value can vary significantly in real-life scenarios since the single bilayered vesicles were created before RNA encapsulation. However, the observed difference in thickness between these two LNPs can be attributed to the choice of ionizable lipids. The bilayer width plays a crucial role in the overall integrity and functionality of the LNPs, impacting their interaction with biological membranes, cellular uptake, and the release of encapsulated cargo, such as mRNA.

We calculated the order parameter of the ionizable lipids in mRNA-1273 and BNT162b and observed a significant difference between the two lipid types. The order parameter quantifies the degree of order in the acyl tails of lipid molecules, providing insights into how well-aligned or disordered these tails are within a membrane environment. Mathematically, it is defined as, 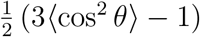 where *θ* is the angle between the vector representing the lipid tail and the normal to the membrane surface. An order parameter value close to 1 indicates high ordering (as seen in a gel phase), where the lipid tails are well-aligned, while a value close to 0 signifies disordered lipid tails (typical of a fluid phase), indicating greater flexibility and less alignment. Negative values suggest that the tails are anti-aligned. As shown in Figure 6, the order parameter values for the ionizable lipids of mRNA-1273 are significantly higher and closer to 1 compared to those of BNT162b. This suggests that the ionizable lipids in mRNA-1273 are more ordered than those in BNT162b. From this we can also understand that the ionizable lipid of BNT162b are more bifurcated compared to the structure of ionizable lipids of mRNA-1273.

**Figure 6:**
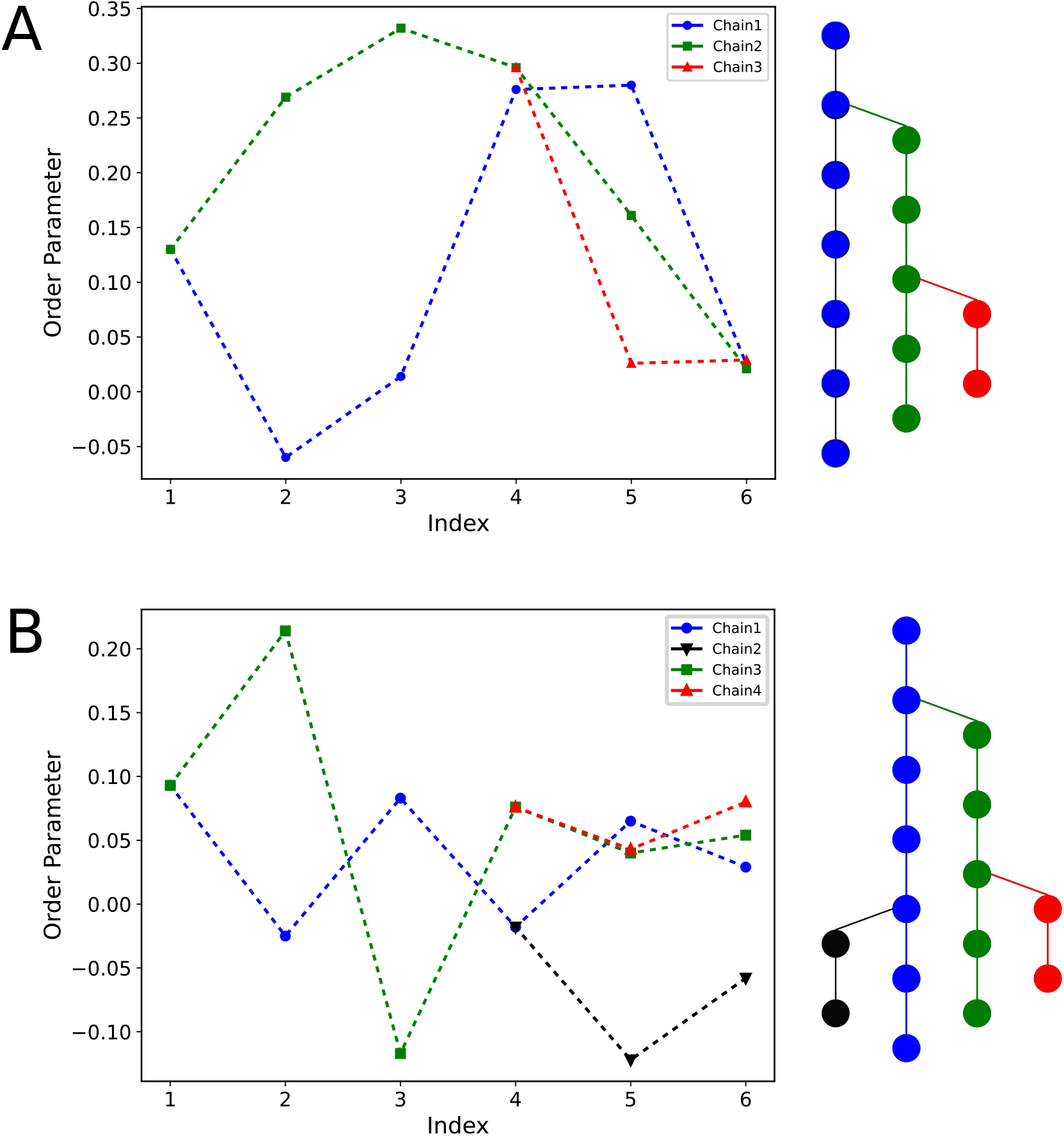
Order parameter of ionizable lipids of (a) mRNA-1273 and (b) BNT162b. The chains of the ionizable lipids corresponding to the order parameters are shown in the right. Index represents the bonds from the first beads of the head group.

To understand any potential aggregation of ionizable and phospholipids, we illustrated the distribution of DPPC and ionizable lipids within the bilayer simulation for both mRNA-1273 and BNT162b, in the figure S9 of supporting information. The analysis reveals that both mRNA-1273 and BNT162b exhibit distinct lipid distributions without any signs of aggregation, suggesting stable interactions within the lipid environment.

## Conclusion

The simulated model for vesicles containing lipids of two important COVID-19 vaccines, mRNA-1273 (Moderna) and BNT162b (Pfizer), were studied. In this study, we used a coarse-grained martini model to create lipid vesicles. It was observed that both the vesicles are stable over a 1500 ns period. As seen from the distribution of lipids, it was observed that due to the presence of a more branched chain in the tail of ionizable lipid of BNT162b (ALC-0315), the lipids are seen as more bifurcated compared to the structure of ionizable lipids of mRNA-1273 (SM-102). In the density distribution of lipids, although both the LNPs have very similar density profile, it is observed that the head group of ionizable lipid, ALC-0315 in BNT162b has a broader profile compared to the SM-102 in mRNA-1273. This suggests that the mRNA-1273 has tighter packing during lipid formation compared to BNT162b. It is shown in earlier studies that BNT162b is stored at a much lower temperature (−80 °C) than mRNA-1273 (−20 °C).^37^ While the impact of this extreme cold on lipid stability isn’t entirely clear, it’s possible that temperature plays a role in lipid behavior.

## Acknowledgement

The authors thank the support and the resources provided by PARAM Sanganak under the National Supercomputing Mission, Government of India at the Indian Institute of Technology, Kanpur. MRB thank Indian Institute of Technology Kanpur for financial support.

