## Supporting Information for "Comparative Analysis of Lipid Nanoparticles in Pfizer-BioNTech and Moderna COVID-19 Vaccines: Insights from Molecular Dynamics Simulations"

### Coarse Grained Mapping for ionizable lipids and PEG molecules

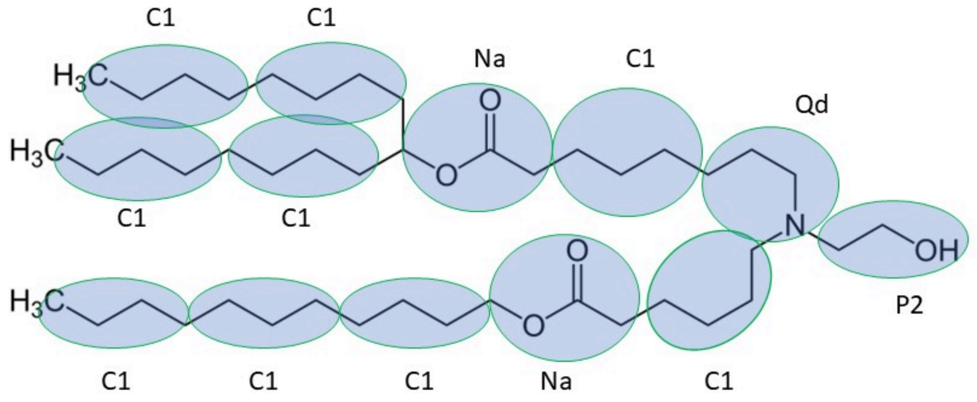


Figure S1: Course grained mapping for ionizable lipid of mRNA-1273 (SM-102). Martini 2 bead types are assigned.

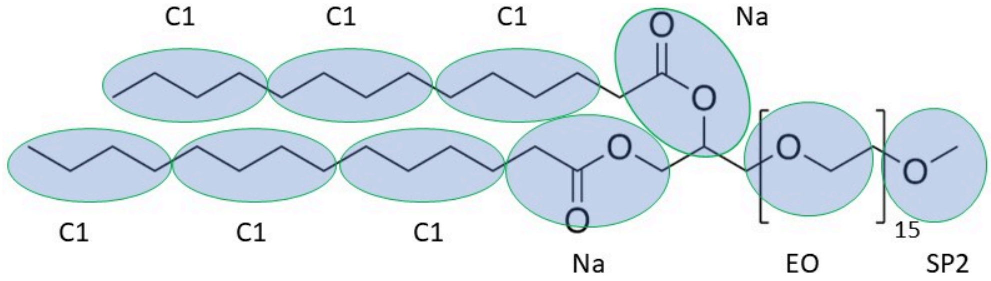


Figure S2: Course grained mapping for ionizable lipid of mRNA-1273 (PEG2000-DMG). Martini 2 bead types are assigned.

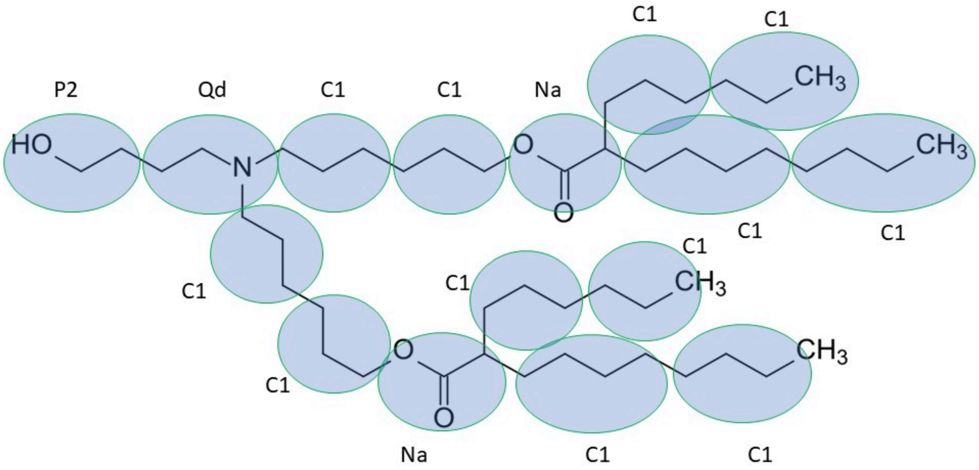


Figure S3: Course grained mapping for ionizable lipid of BNT162b2 (ALC-0315). Martini 2 bead types are assigned.

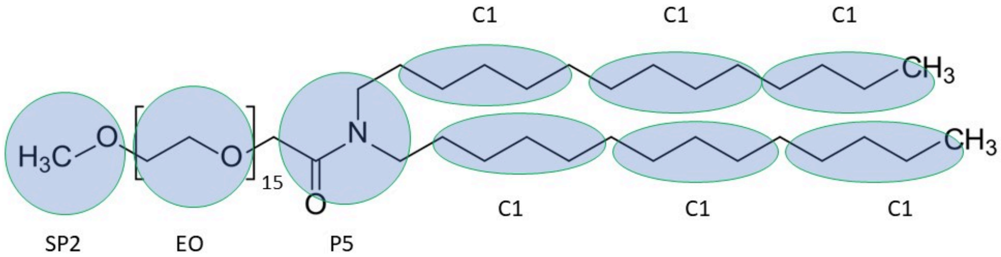


Figure S4: Course grained mapping for PEG lipid of BNT162b2 (ALC-0159). Martini 2 bead types are assigned.

#### Vesicles with mRNA encapsulated after 1500 ns

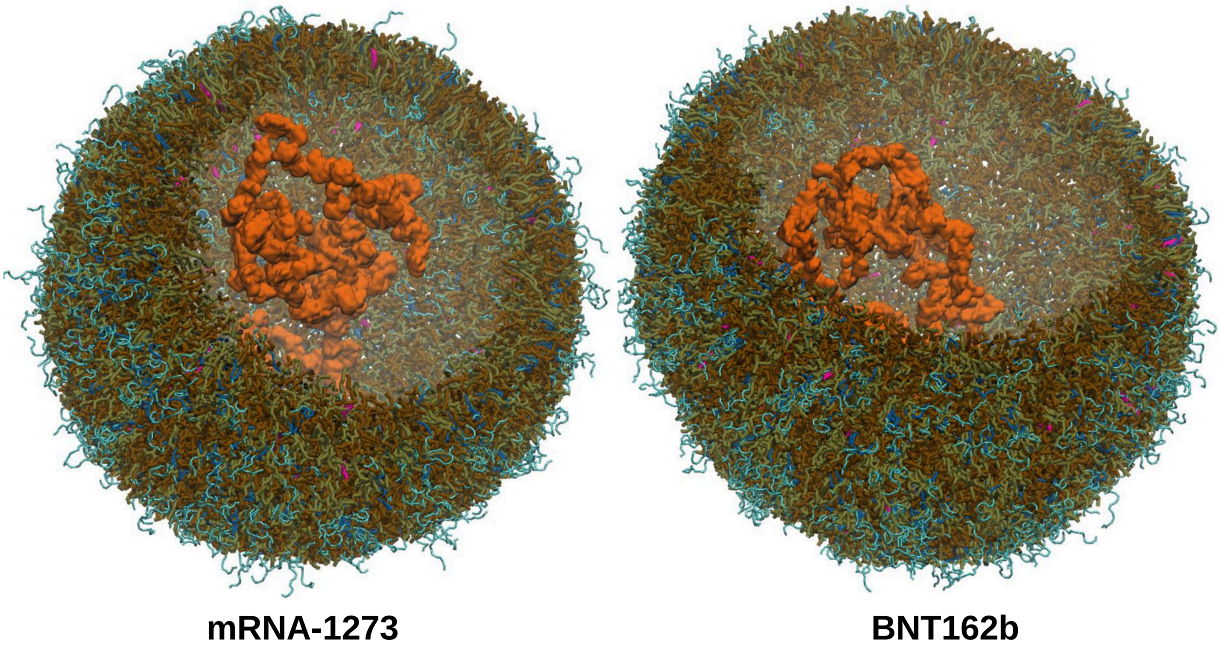


Figure S5: After 1500 ns a stable structure of lipids with mRNA encapsulated were observed in both mRNA-1273 and BNT162b. The water was not shown for better visibility. PEG (cyan), DPPC (brown), ionizable lipids (ochre), cholesterol (magenta) and mRNA (orange).

#### Lipid arrangements in the different layer of vesicles

To understand the arrangement of lipids especially ionizable lipids and phospholipids, we have looked into it in different layer. As our vesicle is speherical, we have taken four different slices as shown in Figure S6.

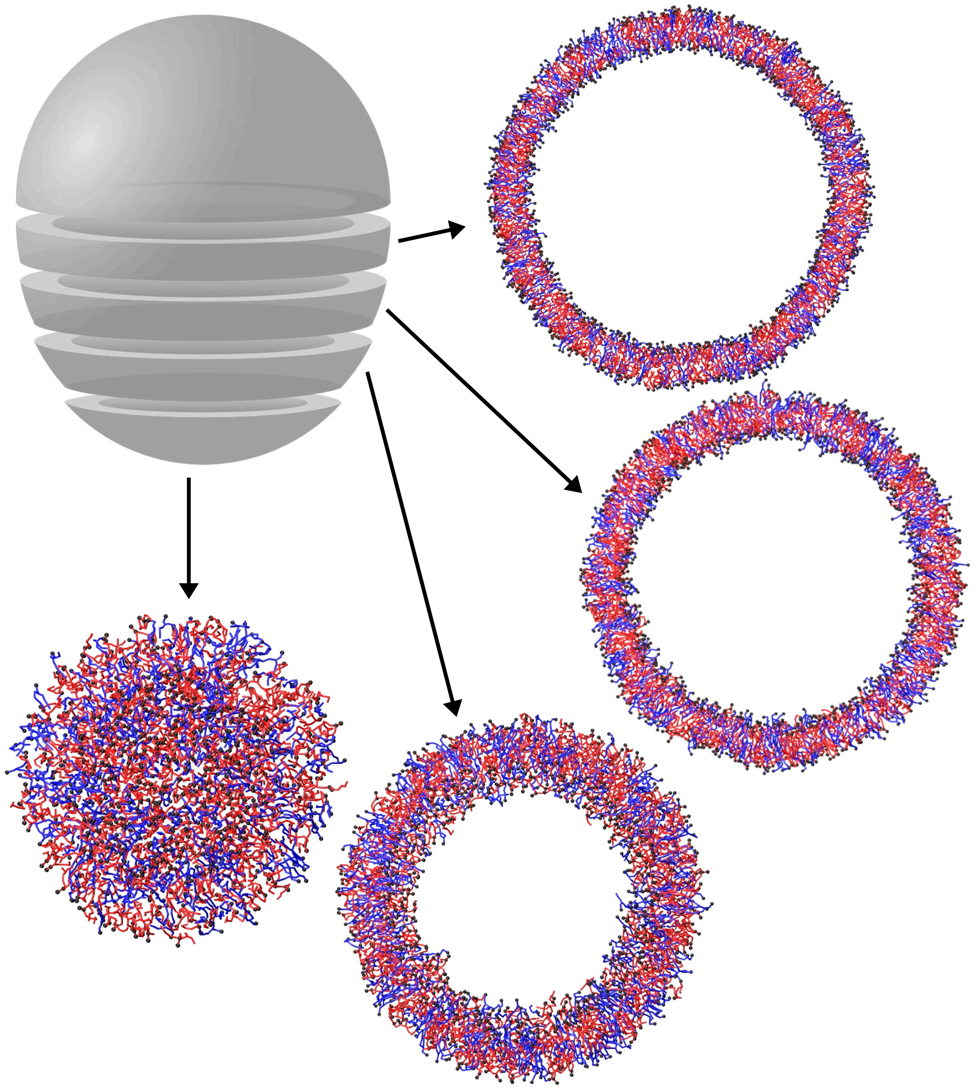

Figure S6: Schematic diagram of different layer of vesicle.

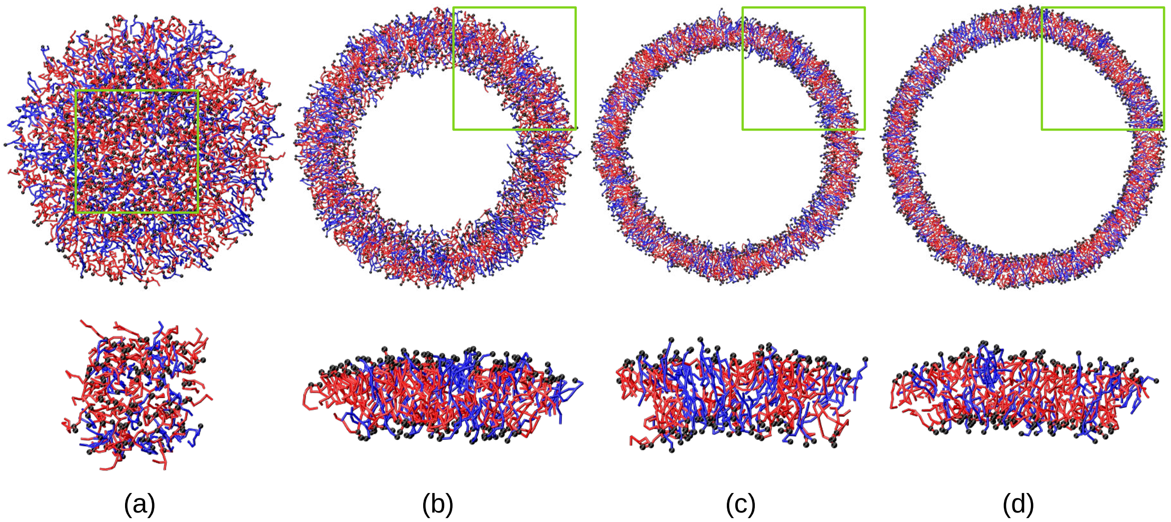


Figure S7: Distribution of DPPC and ionizable lipids of mRNA-1273 at different cross section of the vesicles. Correspond zoomed images are shown in the image below. DPPC is shown in blue and ionizable lipids shown in red.

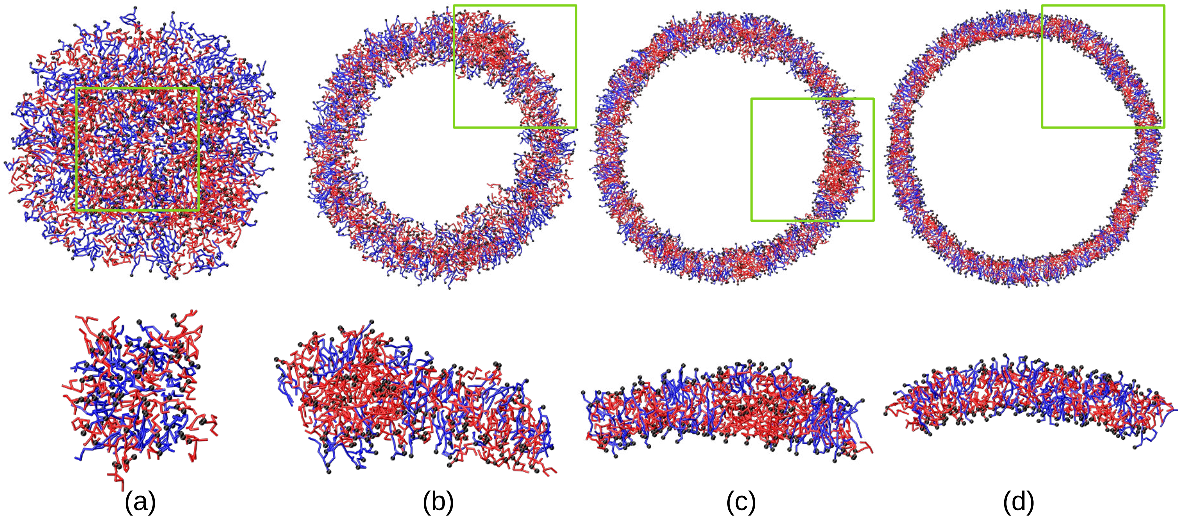


Figure S8: Distribution of DPPC and ionizable lipids of BNT162b at different cross section of the vesicles. Correspond zoomed images are shown in the image below. DPPC is shown in blue and ionizable lipids shown in red.

### Lipid distribution in the bilayer

Before constructing a vesicle, we initially created a lipid bilayer. To analyze the lipid distribution and assess any potential aggregation, the figure below illustrates the arrangement of the lipids. Our observations indicate that there is no aggregation of lipids in either mRNA-1273 or BNT162b.

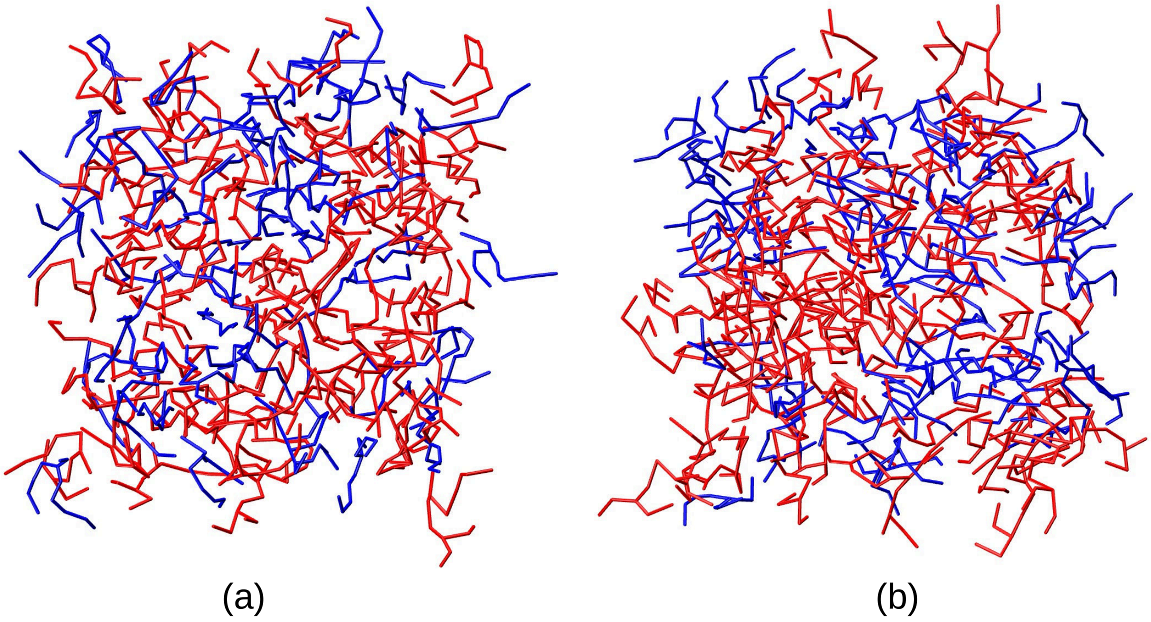


Figure S9: Distribution of DPPC and ionizable lipids in the bilayer simulation for (a) mRNA-1273 and (b) BNT162b. DPPC is represented in blue, while the ionizable lipids are shown in red.
